# Maternal cortisol and emotional complaints alter breast milk’s microbiome composition

**DOI:** 10.1101/2022.09.28.509979

**Authors:** Nadia Deflorin, Ulrike Ehlert, Rita T. Amiel Castro

## Abstract

Breast milk (BM) is considered the “gold standard” of nutrition due to its many benefits for the infant. On the other hand, the challenges of pregnancy can lead to increased stress in some women, which may affect BM quality. Although studies have demonstrated a negative link between maternal psychopathology and child development, it remains unclear how maternal psychobiological changes can be intergenerationally transmitted.

**Aims:** We investigated the associations between maternal biological stress, depressive symptoms, self-reported stress and anxiety symptoms and the BM microbiome. Further, we analysed these parameters in relation to BM glucocorticoid concentrations and explored the influence of BM glucocorticoids on BM bacterial composition.

**Methods:** N=100 women completed standardised questionnaires (e.g. EPDS, STAI, GAS) at 34-36 weeks gestation and in the early postpartum; and sampled saliva at 34-36 and 38 weeks gestation. BM samples were collected in the early postpartum. Microbiota was analysed using 16S rRNA amplicon sequencing.

**Results:** Prenatal GAS and pregnancy-related symptoms (sum) were negatively correlated with *Alphaproteobacteria* (τ = - 0.2137, FDR = 0.0199*; τ = - 0.1805 FDR = 0.0798*), whereas in the postpartum period, the STAI-S scores were negatively correlated with different taxa. Postpartum-related symptoms (sum) were also linked to decreased *Propionibacteriales*. Salivary cortisol AUCg at 34-36 weeks was negatively correlated with *Actinomycetia* (τ = -0.1633, FDR = 0.0630*) and *Clostridia* (τ = -0.1824, FDR = 0.0630*). BM cortisol was negatively correlated with *Staphylococcus* (τ = - 0.2047, FDR = 0.0885*) and positively correlated with BM alpha diversity. No associations emerged between psychobiological parameters and BM glucocorticoids.

**Conclusions:** Increased perinatal emotional complaints and prenatal cortisol AUCg were associated with decreased commensal bacteria, whereas enhanced BM cortisol was linked to increased alpha diversity and reduced *Staphylococcus*. These findings suggest a negative relation between high maternal psychobiological complaints and commensal milk microbiota.

## 1. Introduction

Breast milk (BM) is considered the “gold standard” of nutrition due to its many benefits for the infant, including fewer respiratory infections and a lower incidence of obesity, type 1 and 2 diabetes mellitus, and gastroenteritis [1, 2]. BM composition is dynamic, highly complex and varies within one feeding, through the course of the day, across lactation periods, and between populations [3, 4]. BM has long been considered sterile, yet recent studies have shown it to be a rich source of commensal microbes that can positively influence the child’s health [5]. For instance, in a recent review [6], the BM microbiome was found to be quite diverse, containing more than 800 bacterial species. *Serratia, Ralstonia, Propionibacterium, Pseudomonas, Staphylococcus, Streptococcus, Corynebacterium, Sphingomonas and Bradyrhizobium* represent the nine genera that form the core BM microbiome [5, 7] as they constitute approximately half of its microbial community [8, 9]. Evidence suggests that BM is not solely a passive reflection of maternal bioactive substances, but is rather an actively regulated secretion that may program the infant’s cognitive and immunological development [10, 11]. BM microbial community reaches the child’s gut via breastfeeding, influencing its colonisation [12, 13], and can affect several aspects of newborn health [12, 14–17]. Interestingly, it has recently been demonstrated that BM bacterial, hormonal and immunological composition may be negatively altered by maternal psychopathology [11, 18].

The changes and challenges of pregnancy can lead to increased stress in some women [19–22], which may in turn negatively affect BM quality [18, 23, 24]. Other determinants that have been found to interact with BM composition include maternal age, parity, body mass index (BMI), diet, use of antibiotics, and mode of delivery [5]. Although there is a considerable body of research demonstrating a negative link between perinatal maternal psychopathology and child development and health [11, 18, 25, 26], it remains unclear how maternal psychobiological complaints can be intergenerationally transmitted [27]. One strong candidate in this regard is BM. BM composition can be shaped by multiple factors such as maternal body index and diet [11]. The development of the maternal BM microbiome begins in the third trimester of pregnancy [28, 29], with bacteria from the maternal gut probably entering the mammary gland through bacterial translocation [30–32]. This potential microbiome transfer route is known as the entero-mammary pathway [12, 29, 32] and can be influenced, among others, by maternal emotional complaints [16, 33, 34] via the microbiome-gut-brain axis [35]. The latter describes a bidirectional interaction occurring between the gastrointestinal tract, the gut microbiome, and the central nervous system [35]. Communication pathways of this axis include vagal nerve activation, neurotransmitters, metabolites, as well as the immune system [16]. Consistent with these pathways, experimental studies also suggest a bidirectional communication between the gut microbiome and the hypothalamic-pituitary-adrenal (HPA) axis [36].

The HPA axis is the main system controlling stress responses [37], in which the stress hormone cortisol is produced by the adrenal glands in response to physiological and psychological stress [38, 39]. Cortisol released in the systemic circulation appears to be the primary source of BM glucocorticoids, as BM cortisol was shown to be strongly correlated with plasma cortisol concentrations [25, 40]. Maternal stress is a substantial factor influencing higher cortisol concentrations in BM [24, 41–43]. Although BM glucocorticoids were first detected in the 1970s [44], few studies to date have looked at their effects on the offspring [25].

Given that BM is an underexplored pathway underlying the relation between perinatal emotional complaints and the offspring, the aim of the present study is threefold. First, we investigated the associations between maternal perinatal biological stress, depressive symptoms, self-reported stress, and anxiety symptoms and the BM microbiome. Second, we analysed these parameters in relation to BM glucocorticoid concentrations. Finally, we explored the influence of BM glucocorticoids concentrations on BM bacterial composition. To the best of our knowledge, this is the first study to comprehensively examine the interplay between the BM microbiome and glucocorticoid concentrations in colostrum (the initial BM) in relation to maternal perinatal mental health. This is relevant to improve our understanding of the interactions between maternal emotional complaints, endocrinological parameters and bacterial composition [5, 10, 11]. We hypothesise that women with higher levels of perinatal emotional complaints and biological stress will show lower bacterial diversity and less commensal bacteria as well as elevated BM glucocorticoid concentrations. Additionally, we hypothesise that higher levels of BM cortisol can negatively alter the BM microbiome.

## 2. Materials and Methods

From January 2021 to January 2022, we recruited women in the German-speaking part of Switzerland through online platforms, mailing lists, social media, the study’s homepage as well as flyers handed to patients receiving prenatal medical care at gynaecology practices in and around Zurich and Aargau. To meet the inclusion criteria, women had to be between 18 and 45 years old, currently in the 3^rd^ gestational trimester, fluent in German, of European origin and physically healthy according to self-report. Women were excluded if they had a multifoetal gestation, a pregnancy through assisted reproductive technology, medical complications and/or surgical interventions that might have affected ovarian function prior to pregnancy (e.g., polycystic ovary syndrome) or a planned caesarean section. Additional exclusion criteria were current intake of hormones, psychotropic substances, drugs (e.g.cocaine, marijuana) and antibiotics. A total of *N* = 119 prenatal women and their children were included in this prospective cross-sectional study.

The procedures followed were in accordance with the ethical standards of the Zurich Cantonal Ethics Committee on human experimentation and approval was obtained prior to study commencement. Ethical approval was granted by the Zurich Cantonal Ethics Committee (KEK-ZH-Nr. 2020-01928) and the study was conducted at the Psychological Institute of the University of Zurich in accordance with the Declaration of Helsinki.

The inclusion and exclusion criteria were screened using an online questionnaire. Women who met the inclusion criteria according to this questionnaire then received the study information and consent forms via postal mail. Next, a telephone interview was conducted with interested participants, in which the inclusion criteria were re-checked and the Structured Clinical Interview for DSM-5 (SCID-5-CV) was administered. Eligible participants were invited to the first laboratory appointment between the 34^th^ and 36^th^ week of gestation (T1), conducted either as a home visit or via videotelephony, depending on the respective COVID-19 restrictions at the time. Before T1, participants were asked to sign the consent forms. T1 followed a standardised protocol. First, the procedure was explained and maternal anthropometric measures were recorded. In the case of videotelephony appointments, this information was taken either from the participants’ maternity records or their most recent gynaecological examination. Next, instructions regarding saliva sampling were provided, both verbally and in writing, as participants were asked to independently collect saliva samples during the study. Finally, any potential questions were clarified and participants were asked to complete online questionnaires on socio-demographic, psychological and behavioural information.

On the day after the first appointment (T1), participants were asked to collect saliva and briefly reply to some questions on their perceived stress levels and sleep quality. Saliva sampling took place on two consecutive days immediately after awakening, 30 min and 45 min after awakening, and before going to bed. The second saliva collection was scheduled for the 38^th^ week of pregnancy (T2), again accompanied by the same brief questionnaire as in T1. Participants were asked to freeze the saliva samples in their home freezer. Within the first 120 hours after birth, preferably on the fourth day postpartum, the final study appointment (T3) was conducted between 8:00 and 8:30 in the morning as a home or clinic visit. During this visit, any remaining questions were clarified, the procedure was explained, and BM collection was conducted. For the planned analyses, 5-15 millilitres of BM were collected using an electric BM pump. At T3, we also collected the saliva samples that had been stored in the participants’ homes. Subsequently, participants were asked to complete questionnaires related to birth information and some of the psychological parameters from T1.

### 2.1 Instruments

#### 2.1.1 Maternal emotional complaints

Validated psychometric questionnaires were used to assess perinatal depressive symptoms, anxiety symptoms, and self-reported stress.

We measured stress using the German version of the *Perceived Stress Scale* (PSS-10) at T1 and T3 [45]. This is a self-report scale measuring the degree to which the respondent perceives situations to be unpredictable, uncontrollable, and stressful referring to the past four weeks [46]. The 10 items are rated on a five-point Likert scale (0 = “*never*” to 4 = “*very often*”), with higher scores indicating a higher level of perceived stress [45]. Analyses were conducted with the total score as a continuous variable. Cronbach’s alpha in the present study ranged between 0.823 (T3) and 0.828 (T1).

We used the German version of the *General Health Questionnaire* (GHQ-12) [47], which is a reliable instrument to screen for psychological distress [48, 49]. The 12 items are rated on a four-point Likert scale (0 = “*no, not at all*” to 4 = *“much harder than usual*”) [50]. Higher scores indicate higher levels of mental distress [48]. In the present analyses, we used the GHQ-12 total score as a continuous variable. Cronbach’s alpha for this study ranged between 0.780 (T3) and 0.808 (T1).

Pregnancy-specific stress at T1 was measured using the German version of the revised *Prenatal Distress Questionnaire* (NuPDQ) [51]. Respondents are asked to indicate the extent to which they feel “bothered, upset, or worried” about issues that become more relevant as pregnancy progresses (e.g., concerns about caring for a newborn) [52], with items rated on a three-point Likert scale (from 0 = “*not at all*” to 2 = “*very much*”). In the present study, the total score was used as a continuous variable for statistical analyses. Cronbach’s alpha was .073.

The German version of the *Edinburgh Postnatal Depression Scale* (EPDS) was applied to assess maternal depressive symptoms at T1 and T3 [53]. The questionnaire was developed to assess depressive symptoms in pre- and postnatal women. It consists of 10 items rated on a four-point Likert scale (0 = “*yes, most of the time*” to 3 = “*no, never*”) relating to how one has felt in the last seven days. In our analyses, we used the total score as a continuous variable. Cronbach’s alpha for this study ranged between 0.798 (T1) and 0.880 (T3).

Pregnancy/birth anxiety was assessed at T1 using the *Birth Anxiety Scale* (Geburts-Angst-Skala, GAS) [54]. The scale comprises 25 items assessing the extent to which respondents are afraid of certain aspects related to pregnancy and birth, rated on a four-point Likert scale (0 = “no fear” to 3 = “strong fear”) [55]. For the present analyses, the total score was used as a continuous variable. Cronbach’s alpha was 0.871.

Trait anxiety and state anxiety were measured using the German version of the State-Trait Anxiety Inventory (STAI) [56, 57]. The State scale (STAI-S) addresses the level of anxiety in a particular situation [56] and consists of 20 items rated on a four-point Likert scale (1 = “*not at all*” to 4 = “*very much*”), with higher scores indicating higher state anxiety [58]. Cronbach’s alpha for this scale ranged between 0.891 (T1) and 0.924 (T3). The Trait scale (STAI-T) relates to the overall disposition to experience worries, fears, and anxieties in different situations [59]. It likewise consists of 20 items rated on four-point Likert scale (1 = “*almost never*” to 4 = “*almost always*”), and asks respondents how they feel in general. Higher scores indicate higher trait anxiety. Cronbach’s alpha for the current study ranged between 0.890 (T3) and 0.897 (T1). Trait and State anxiety total scores were scored separately and treated as continuous variables in our analyses.

Maternal emotional complaints can be considered as a multidimensional concept, with women reporting various different symptoms (e.g., anxiety, stress, and/or depression) [18, 60, 61]. Therefore, we created summary variables from the questionnaires described above. To capture pregnancy-specific emotional complaints, the questionnaires NuPDQ, GAS, and EPDS were merged, as they all specifically target the prenatal period. To do so, we used the total scores from each questionnaire at T1 and summed them together to create a pregnancy-related sum score (continuous variable). Similarly, a multidimensional summary variable was created to capture postpartum emotional complaints. Total scores derived from the GHQ-12, PSS-10, EPDS, and STAI-S were combined into a postpartum sum score (continuous variable), as they account for depressive symptoms, anxiety symptoms and self-reported distress.

#### 2.1.2 Maternal prenatal salivary cortisol

During the course of the study, participants were required to collect saliva samples for the assessment of cortisol concentrations. Under standardised conditions, participants were required to collect 8 saliva samples per time point (T1 and T2), corresponding to a total of 16 saliva samples per participant. The three morning and one evening samples were collected using the passive drool method in SaliCap sampling tubes with 2 mL capacity (SaliCaps, IBL international GmBH, Hamburg, Germany). For this purpose, the women were instructed to collect saliva immediately after awakening, 30 min after awakening, 45 min after awakening and before going to bed. After sampling, they were instructed to store the samples in their freezers until the home or clinic visit (T3). The day before sampling, participants were asked to avoid exercise, alcohol, coffee, black tea, and chewing gum. Additionally, they were asked not to eat, smoke, or brush their teeth during the morning saliva collection sessions. After completion of study participation, samples were transported by a member of the study team using a cool box and stored at -20 degrees in the biochemical laboratory of the University of Zurich. For subsequent cortisol analyses, samples were then thawed, centrifuged, and analysed biochemically using an enzyme-linked immunosorbent assay (ELISA; IBL international GmBH, Hamburg, Germany). Individual values beyond the range of the kit were excluded from the analyses (*N* = 2).

#### 2.1.3 Breast milk collection

Within the first 120 hours after birth, preferably on the fourth day postpartum, the final study appointment (T3) was performed between 8:00 and 8:30 in the morning as a home or clinic visit, as BM cortisol and cortisone concentrations follow the diurnal rhythm of the HPA axis [62]. The method of BM collection was similar to that used by McGuire et al. (2017) [63]. Accordingly, participants were asked to wash their hands and dry them with a disposable towel provided by the study team. Both the participant and the study team member had to disinfect their hands and put on disposable gloves. As the researcher had to unwrap the materials and give them to the woman, this procedure ensured that the materials were as sterile as possible. Once the appropriate BM pump set was selected, the breast was then stimulated. Following this, the breast was cleaned twice with a newly opened pre-packaged castile soap towelette (Pdi Inc. Pyd41900 Castile Soap Towelettes). We used an electric BM pump (Medela Symphony) and disposable pump sets (Medela Inc.) to collect milk samples into a single-use, sterile, 35 ml milk collection container (Medela Inc.). To ensure the best possible milk flow, the breast chosen for pumping was the one that was not used during the last feeding or that the participant felt to be fuller. If not enough milk was collected from one breast, participants were asked to express milk from both breasts; in these cases, the previously described cleaning procedure was repeated with the other breast.

For the BM microbiome sample, we collected 5-10 ml of BM in a milk collection container. In a second container, a further 2-5 ml were expressed for the analyses of BM glucocorticoids. Immediately after collection, for the endocrinological analyses, BM was transferred from the 35 ml milk collection container into 2 ml microtubes (Sarstedt AG) by a member of the study team using a sterile pipette. All samples were immediately placed in a cool box to be transported by members of the research team to the biochemical laboratory at the University of Zurich, where they were stored at -80 degrees until shipment. *N* = 97 BM samples were collected, which amounted to 1.64 litres of BM. Samples for microbiome analysis were shipped to the Clinical Microbiomics Laboratory in Copenhagen (Denmark). Samples for cortisol and cortisone measurements were shipped to the Laboratory of Endocrinology, University Medical Centers (UMC), Amsterdam (Netherlands).

### 2.2 BM cortisol and cortisone concentrations

To evaluate cortisol and cortisone concentrations in BM, isotope dilution liquid chromatography-tandem mass spectrometry (LC-MS/MS) was used, as previously described in Van der Voorn et al. (2015) [64]. Briefly, milk samples were washed with hexane after internal standards (13C3-labelled cortisol and cortisone) were added to the samples. Samples were then extracted with isolute plates (Biotage, Uppsala, Sweden) and analysed by LC-MS/MS (Acquity with Quattro Premier XE, Milford MA, USA, Waters Corporation). The intra-assay coefficient of variation (CV) was 4-5% for cortisol and 5 % for cortisone at different values. The inter-assay coefficient of variation for both, cortisol and cortisone was < 9%, with the lower limit of quantitation of 0.5 nmol/L.

### 2.3 BM 16S ribosomal RNA (rRNA) sequencing and quality control

At the Clinical Microbiomics laboratory, DNA was extracted from ∼5 mL aliquots of the *N* = 97 BM samples. Prior to DNA extraction, cells from the aliquots were pelleted by centrifugation (2 × 10 minutes at 6000 rpm). In a next step, DNA extraction from the pellets was performed using the NucleoSpin 96 Soil (Macherey-Nagel) kit. Bead beating was performed horizontally on a Vortex-Genie 2 at 2700 rpm for 5 minutes. Negative and positive (mock) controls were incorporated into the analysis. Products from nested PCR were pooled based on band intensity, before the resulting library was cleaned with magnetic beads. Pooled library DNA concentration was measured fluorometrically. Sequencing was performed on an Illumina MiSeq desktop sequencer using the MiSeq Reagent Kit V3 (Illumina) for 2x 300 bp paired-end sequencing. We performed PCR using the forward primer S-D-Bact-0341-b-S-17 and the reverse primer S-D-Bact-0785-a-A-21 [65] with Illumina adapters attached. In fact, these are universal bacterial 16S rDNA primers targeting the V3-V4 region. The following PCR program was applied: 98 °C for 30 sec, 29x (98° C for 10 s, 55 °C for 20 s, 72 °C for 20 s), 72°C for 5 min. Amplification was verified by running the products on an agarose gel. Indices were added in a subsequent PCR, utilising an Illumina Nextera kit with the subsequent PCR program: 98 °C for 30 sec, 8x (98° C for 10 s, 55 °C for 20 s, 72 °C for 20 s), 72 °C for 5 min. Attachment of the indices was validated by testing the products on an agarose gel. To account for variations in detection ability due to differences in sequencing depth, all samples were rarefied to a fixed number of reads. Rarefied count and abundance matrices were generated by random sampling of reads without replacement with a target number of 4063 read counts. Any samples with a lower number of reads were excluded.

### 2.4 Statistical analysis

To define the minimum sample size required to test the study hypothesis, an a priori power analysis was performed with G*Power version 3.1.9.7 [66]. All statistical analyses were conducted with the IBM Statistical Package for the Social Sciences (SPSS version 28 for Windows) and the software program R, using the adonis2 function from the vegan R package. Normality of the sample was tested with the Shapiro-Wilk test, and non-parametric tests were used for non-normally distributed data. For associative analysis with salivary cortisol concentrations, we used the mean value of the area under the curve with respect to the ground (AUCg) of the two consecutive days of saliva sampling. The AUCg was itself calculated using the trapezoid formula, which is routinely calculated to include multiple time points [67]. Moreover, we used a mean value of total cortisol decline of the two consecutive days, calculated as follows: Δ in the morning - Δ during the day, in which Δ in the morning corresponds to the highest morning value - baseline morning value and Δ during the day corresponds to the highest morning value - the evening value. Both procedures, the AUCg and the total decline, were conducted separately for T1 and T2. Spearman’s rank correlation was used to examine correlations between and within psychological variables (PSS-10, GHQ-12, NuPDQ, EPDS, GAS, STAI-S, pregnancy-related sum, postpartum sum), salivary cortisol and BM glucocorticoid concentrations at all time points. Additionally, a Kruskal-Wallis test was run to test for significant differences in BM glucocorticoid concentrations depending on the extraction breast (e.g. right, left or both). Next, the cortisone-to-(cortisol + cortisone) ratio of glucocorticoids in breast milk was calculated [62, 68, 69] in order to describe the activity of 11β-hydroxysteroid dehydrogenase type 2 (11β-HSD2) enzyme, which converts active cortisol into inactive cortisone [68, 69]. The following formula was used for this purpose: BM cortisone/BM cortisone + BM cortisol. Associations between the cortisone-to-(cortisol + cortisone) ratio and the biological or psychological stress markers were evaluated using Spearman’s rank correlation. Both BM glucocorticoids and salivary cortisol results were reported in nmol/l.

For the analysis of the BM microbiome, BM composition was first summarised in an amplicon sequence variant (ASV) abundance table using a customised pipeline based on dada2 [70]. The taxonomic classification of the detected ASVs was performed in two steps. First, ASV sequences were compared with full-length 16S sequences in an internal reference database (CM_16S_27Fto1492R_v1.0.0) using a naive Bayesian classifier. Then, we optimised the taxonomic assignments based on the exact sequence identity percentages between the discovered ASVs and the reference amplicons in an internal V3V4 amplicon database (CM_16S_341Fto785R_v1.0.0.rds). Subsequently, we explored the diversity of the BM microbiome communities within and between samples. For this purpose, bacterial alpha diversity of the samples was calculated using the number of detected entities (richness) and the Shannon index. Bray-Curtis dissimilarity and weighted UniFrac distances were used to calculate bacterial beta diversity. Both alpha and beta diversity estimates were calculated from rarefied abundance matrices. Additionally, when conducting statistical analyses, the Benjamini-Hochberg (BH) method was applied to control for the false discovery rate (FDR) at a level of 10% to control for multiple testing [71].

Associations for beta diversity – including Bray-Curtis dissimilarity and weighted UniFrac distances as dependent variables and psychological variables as independent variables – were performed using a distance-based redundancy analysis (dbRDA) test conducted with 1000 permutations. We ran associative analyses with alpha diversity and taxon abundance as dependent variables and psychological continuous variables as independent variables using Kendall rank correlations. Unrarefied data were used for the associations involving abundance of microbial taxons, for which 97 BM samples were included. For the analyses of alpha and beta diversity parameters, we used 96 BM samples analysed with rarefied data. This difference is due to the rarefaction method (see 2.5, BM sequencing and quality control). Finally, to investigate the potential influence of several covariates on the BM microbiome, PERMANOVA (for categorical variables) and dbRDA (for continuous variables) were performed, following the approach suggested by Falony et al., 2016 [72]. Only covariates that demonstrated a significant relationship with the BM microbiome were further analysed for their predictive effects.

#### 2.4.1 Covariates

Maternal age, parity, maternal BMI, the newborn’s weight and length, and number of hours between last feeding and BM collection were analysed as continuous variables. Categorical variables were education (coded as 1 = university degree or higher and 0 = other) and mode of delivery, considering membrane rupture during labour (coded as 0 = no and 1 = yes). All covariates were identified based on previous literature [5, 12, 63, 73–76].

## 3. Results

### 3.1 Sample characteristics

Descriptive characteristics of the sample are shown in Table 1. Most of the participants were married (66%) and highly educated, with 67% having a university degree. 71% of the women experienced a natural vaginal birth whereas 16% underwent an emergency caesarean section. Of those who had a caesarean section, 43.75% (*N* = 7) had membrane rupture prior to the intervention. The median time of BM sample collection was 97 hours after birth (IQR = 30).

**Table 1.**
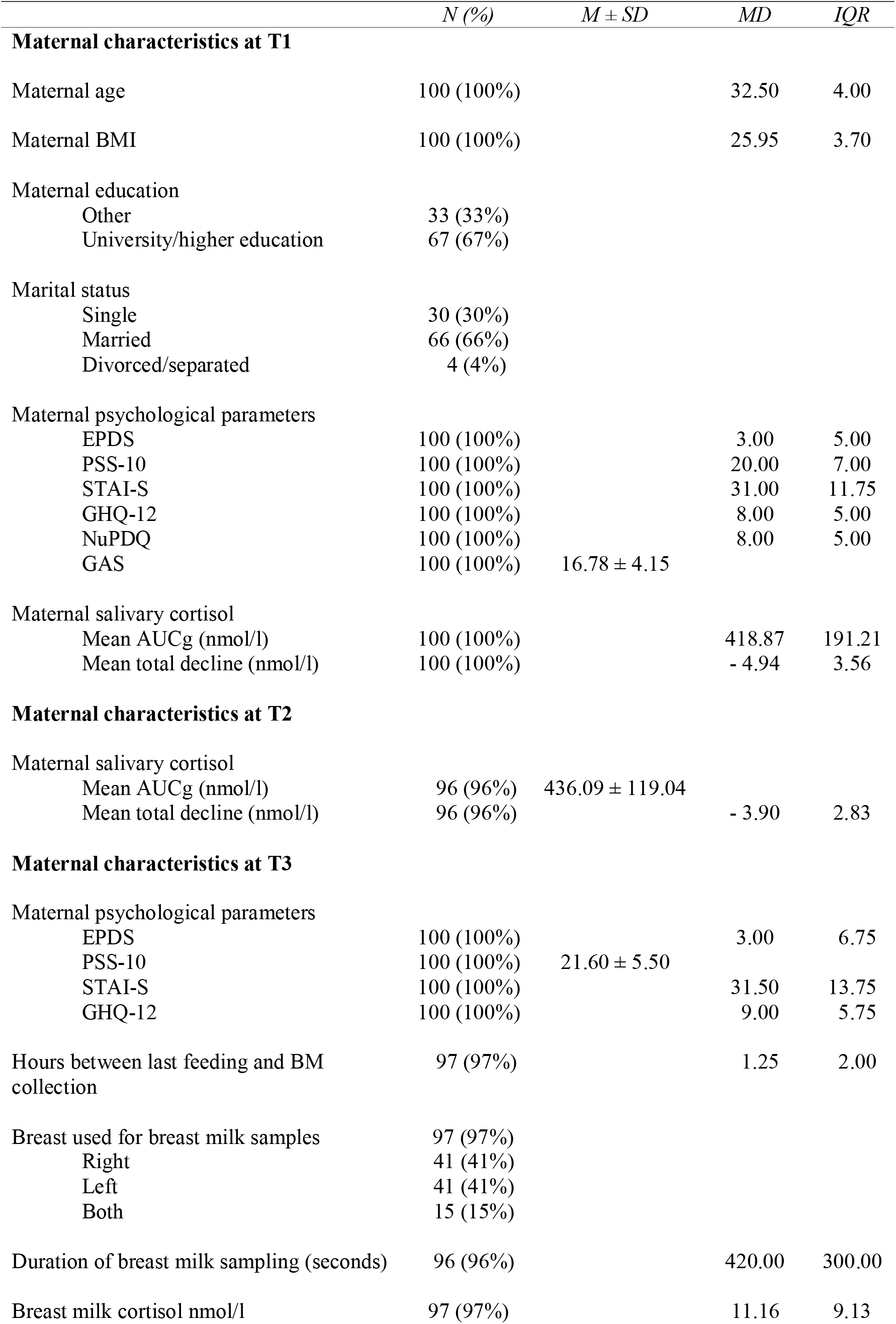
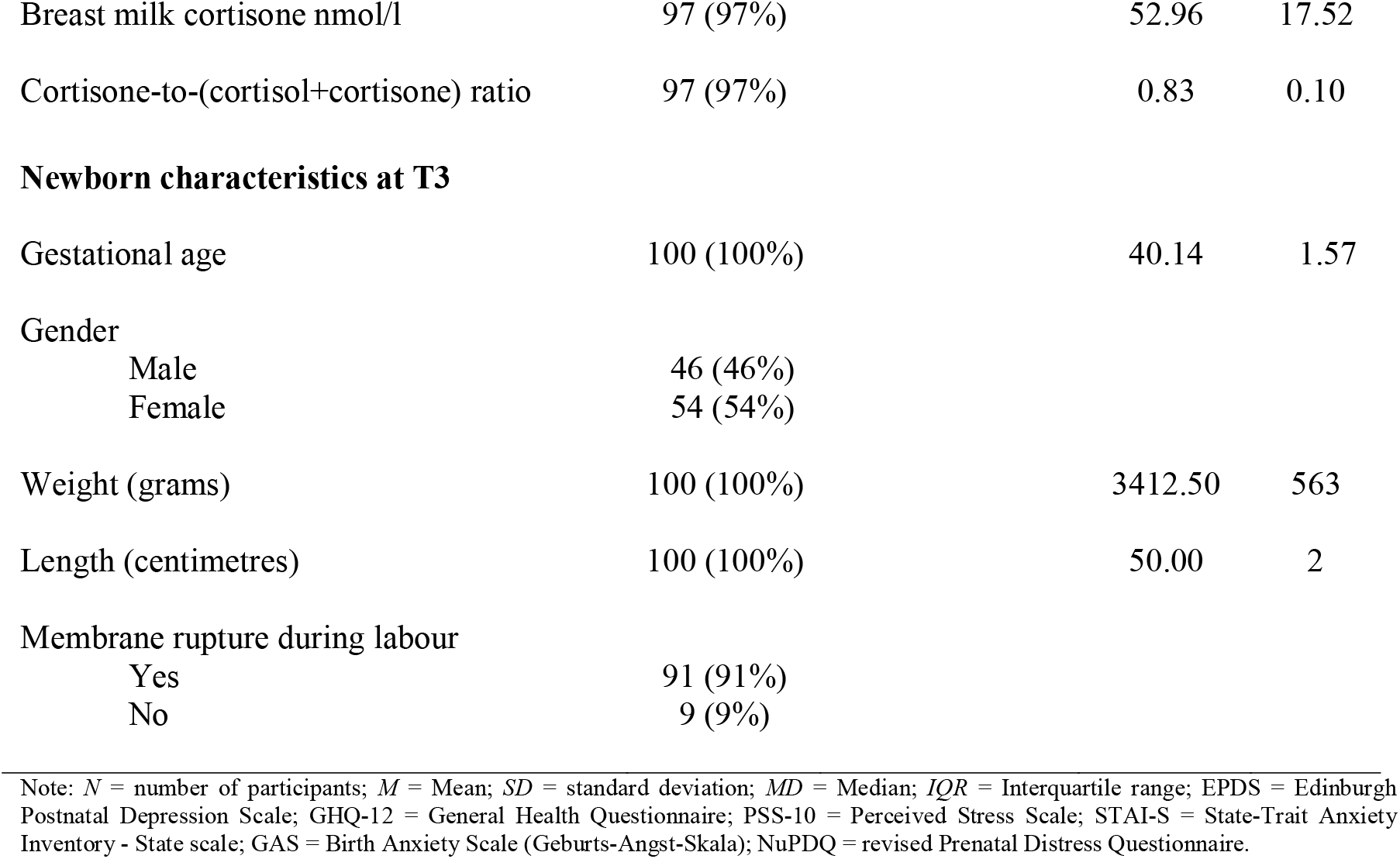
Demographic and obstetric outcomes of the total study sample (*N* = 100)

*N* = 19 participants voluntarily withdrew or were excluded from the study (e.g. pregnancy complications), resulting in a final sample of 100 pregnant women (see Figure 1) with an age range from 22 to 40 years (*MD* = 32.50, *IQR* = 4.00). Our power analysis [66] suggested a required sample size of *N* = 84 to reach 80% power to detect a medium effect (f = 0.15), at a significance level of α = .05. Thus, the achieved sample size of *N* = 100 is sufficient to test the study hypothesis. Moreover, we were unable to collect BM samples from *N* = 3 participants, who had little or no milk (N=2) or were prevented from attending the final appointment due to problems with the newborn baby (N=1).

**Figure 1.**
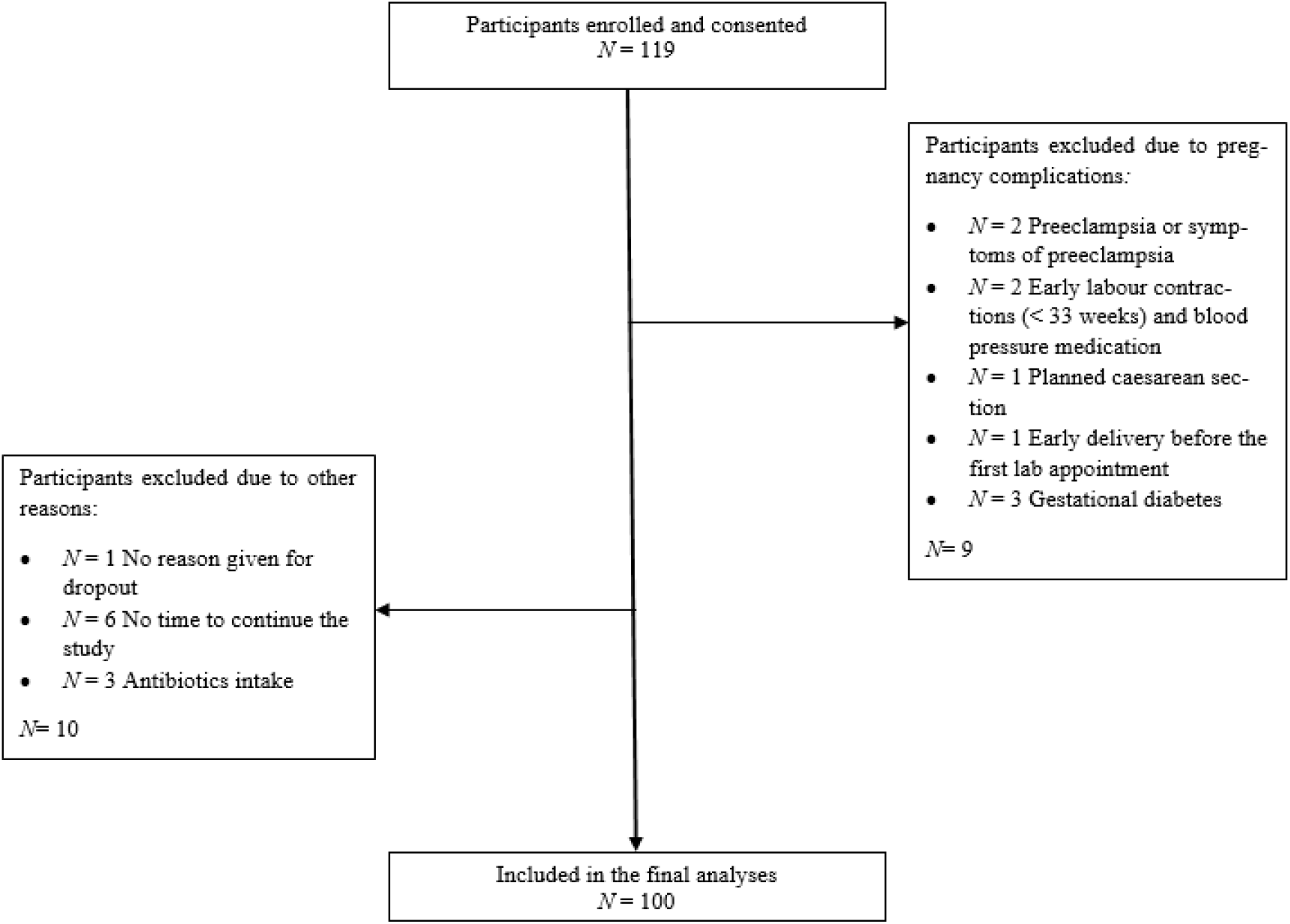
Flow diagram for study participants.

### 3.2 Analysis of the BM microbiome and BM glucocorticoids

For a taxonomic overview, Figure 1 and Figure 2 show the relative abundance of bacterial taxa aggregated at the genus and family levels in the BM. We found that the BM microbiome is dominated by the genera *Staphylococcus* (41.17%), *Streptococcus* (35.26%) and *Gemella* (5.74%) (Figure 1). At the family level (Figure 2), we found that most of the assigned BM taxa were from *Staphylococcaceae* (41.17%), *Streptococcaceae* (35.38%) and *Gemellaceae* (5.74%). Both *Staphylococcus* as well as *Streptococcus* belong to the previously described core BM microbiome [5–7]. Overall, the 97 BM samples had an average of 107.5 thousand (K) read pairs per sample and a minimum of 0.3 K read pairs. An average of 52.4 K read pairs per sample could be mapped to the ASV catalogue, representing an average of 48.1% of the high-quality reads.

**Figure 2.**
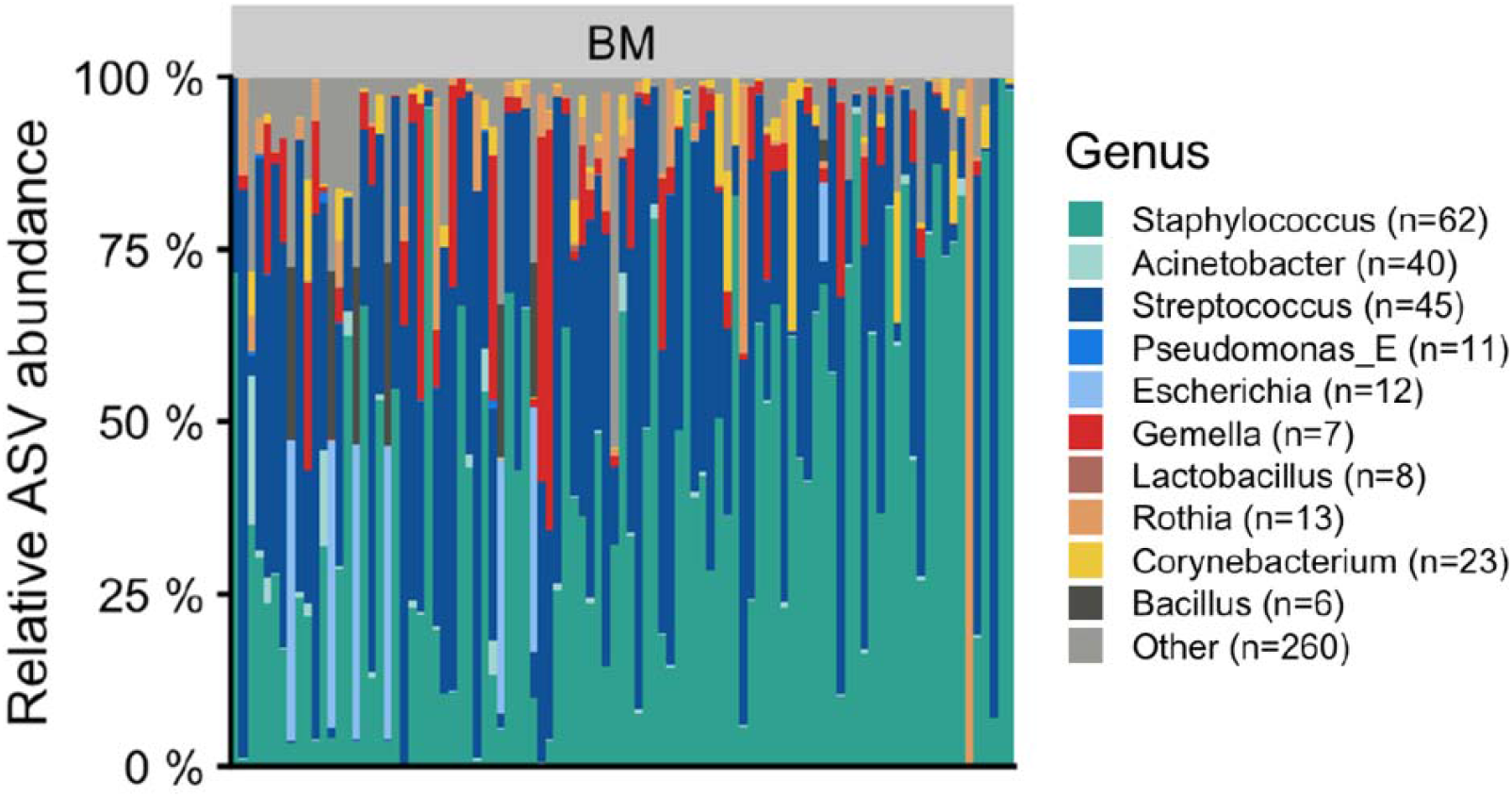
Taxonomic overview of the breast milk at genus level per sample for unrarefied data. The numbers in parentheses refer to the number of ASVs aggregated for a given taxon. Bar plots display the relative abundance of the top 10 taxa with the highest average abundance across all samples. Light grey (Other) indicates the total relative abundance of ASVs that are not in the top 10 most abundant taxa; BM= breast milk.

**Figure 3.**
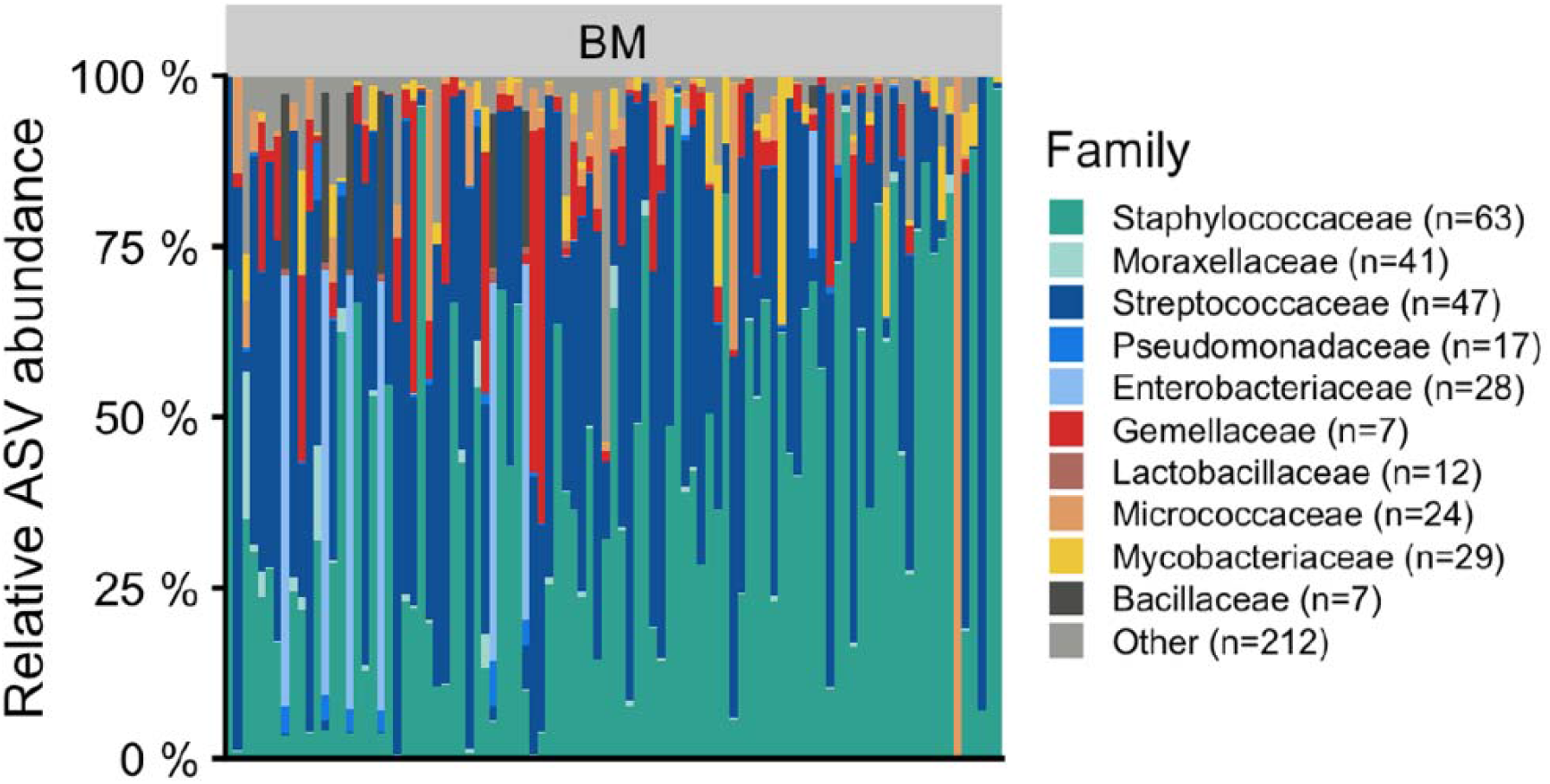
Taxonomic overview of the breast milk at family level per sample for unrarefied data. The numbers in parentheses refer to the number of ASVs aggregated for a given taxon. Bar plots display the relative abundance of the top 10 taxa with the highest average abundance across all samples. Light grey (Other) indicates the total relative abundance of ASVs that are not in the top 10 most abundant taxa; BM= breast milk.

At postpartum, a total of 423.5 millilitres of BM for glucocorticoid analyses were obtained from all 97 participants. The median BM cortisol concentration was 11.16 nmol/l (IQR = 9.12) whereas the median BM cortisone concentration was 52.96 nmol/l (IQR = 17.53). Furthermore, a significant positive correlation between these two concentrations was found (rho = .283**, *p* = .005). Interestingly, there were no significant differences in BM cortisol (*H* = 5.376, *p* = .068) and cortisone (*H* = 1.297, *p* = .532) concentrations based on which breast was selected for sampling. The median of the BM cortisone-to-(cortisol + cortisone) ratio was 0.803 nmol/l with an IQR of 0.100.

### 3.3 Association of maternal emotional complaints with the BM microbiome

In our initial analyses, no significant associations were found between any of the covariates and BM microbial composition (*p* > 0.05). Therefore, no covariates were included in the associative analyses. We explored BM bacterial composition in relation to perinatal maternal emotional complaints from class to species level. After applying the Benjamini-Hochberg (BH) method to control for the FDR [71] at a level of 10%, we found that prenatal GAS and pregnancy-related sum were significantly negatively correlated with the class of *Alphaproteobacteria* (τ = - 0.2137, FDR = 0.0199*; τ = - 0.1805 FDR = 0.0798*). Moreover, in the postpartum period (T3), the class *Alphaproteobacteria*, the order *Propionibacteriales*, as well as the genera *Cutibacterium* and *Pseudomonas*_N were negatively correlated only with the STAI-S values and not with the STAI-T (see Table 2). However, no significant associations were identified between prenatal and postnatal maternal emotional complaints and BM alpha (Shannon diversity index range: τ = - 0.0671, FDR = 0.359 to τ = 0.0006, FDR = 0.994) or beta diversity (Bray-Curtis dissimilarity range: Sum of Squares = 0.3812, Pr(>F) = 0.295 to Sum of Squares = 0.1757, Pr(>F) = 0.999; weighted UniFrac distances range: Sum of Squares = 0.1808, Pr(>F) = 0.305 to Sum of Squares = 0.0925, Pr(>F) = 0.903).

**Table 2.** BM microbiome taxa in relation to maternal prenatal emotional complaints (*N* = 97)

| Category | Item | Variable (T1) | Kendall's tau | $p$-value | FDR |
| --- | --- | --- | --- | --- | --- |
| Class | Alphaproteobacteria | GAS | -0.2137 | 0.0028 | 0.0199* |
| Class | Alphaproteobacteria | Pregnancy-related sum | -0.1805 | 0.0114 | 0.0798* |
| Category | Item | Variable (T3) | Kendall's tau | $p$-value | FDR |
| Class | Alphaproteobacteria | STAI-S | -0.1909 | 0.0079 | 0.0551* |
| Order | Propionibacteriales | STAI-S | -0.2413 | 0.0009 | 0.0173* |
| Order | Propionibacteriales | Postpartum sum | -0.2154 | 0.0028 | 0.0541* |
| Family | Propionibacteriaceae | STAI-S | -0.2413 | 0.0009 | 0.0273* |
| Family | Propionibacteriaceae | Postpartum sum | -0.2154 | 0.0028 | 0.0855* |
| Genus | Cutibacterium | STAI-S | -0.2413 | 0.0009 | 0.0419* |
| Genus | Pseudomonas_N | STAI-S | -0.2322 | 0.0019 | 0.0444* |
| Species | Cutibacterium acnes | STAI-S | -0.2488 | 0.0006 | 0.0497* |
| Species | Pseudomonas_N<br>sp003640395 | STAI-S | -0.2322 | 0.0019 | 0.0742* |

Note: $p$ -value = statistical significance, \* $p < .05$ ; FDR = FDR-adjusted $p$ -value; GAS = Birth Anxiety Scale (Geburts-Angst-Skala); STAI-S = State-Trait Anxiety Inventory - State scale; Postpartum sum = GHQ-12, PSS-10, EPDS, STAI-S at T3; Pregnancy-related sum = NuPDQ, GAS and EPDS at T1

### 3.4 Associations between salivary cortisol concentrations and the BM microbiome

We examined the associations of salivary cortisol concentrations at T1 and T2, measured by AUCg and total decline, with each of the taxa found (from class to species level). Again, the Benjamini-Hochberg (BH) method was applied to control for the FDR [71]. Among others, AUCg at T1 was negatively correlated with *Actinomycetia* (τ = -0.1633, FDR = 0.0630*) and *Clostridia* (τ = -0.1824, FDR = 0.0630*). No associations were found between total decline and any bacterial taxa (*p* > 0.05). Moreover, no significant associations emerged between BM alpha and beta diversity in relation to salivary cortisol concentrations (*p* > 0.05). All significant results of the BM microbiome in relation to salivary cortisol concentrations can be seen in Table 3.

**Table 3.** BM microbiome taxa in relation to maternal salivary cortisol concentrations (*N* = 96)

| Category | Item | Variable (T1) | Kendall's tau | $p$-value | FDR |
| --- | --- | --- | --- | --- | --- |
| Class | Actinomycetia | AUCg | -0.1633 | 0.0180 | 0.0630* |
| Class | Clostridia | AUCg | -0.1824 | 0.0157 | 0.0630* |
| Genus | Stenotrophomonas | AUCg | -0.2433 | 0.0010 | 0.0481* |
| Species | Stenotrophomonas<br>bentonitica_A | AUCg | -0.2557 | 0.0007 | 0.0526* |
| Category | Item | Variable (T2) | Kendall's tau | $p$-value | FDR |
| Genus | Kocuria | AUCg | -0.2720 | 0.0012 | 0.0570* |
| Species | Kocuria rhizophila | AUCg | -0.2720 | 0.0012 | 0.0941* |

Note: $p$ -value = statistical significance, \* $p < .05$ ; FDR = FDR-adjusted $p$ -value; AUCg = area under the curve with respect to the ground.

### 3.5 Associations of maternal emotional complaints and biological stress with BM glucocorticoids

We found significant correlations between and within T1 and T3 for the PSS-10, GHQ-12, EPDS, and STAI-S (*p* < .001**). Spearman’s rho analyses between the PSS-10, GHQ-12, NuPDQ, EPDS, GAS, STAI-S, pregnancy-related sum at T1 and BM cortisol and cortisone concentrations yielded null findings (*p* > .05). Similarly, no significant associations were found between BM glucocorticoids and the GHQ-12, PSS-10, EPDS, STAI-S, and postpartum sum at T3 (*p* > .05).

Spearman’s rho correlations between T1 and T2 for each salivary cortisol measurement (maternal salivary cortisol AUCg: rho = .706**, *p* < .001; total decline: rho = .507**, *p* < .001) were significant. However, both salivary cortisol AUCg and total decline at T1 and T2 were not significantly associated with any psychological parameters at T1 and T3 (*p* > .05). Similarly, no significant results were found between the maternal salivary cortisol AUCg and total decline measurements and BM glucocorticoids (*p* > .05).

The BM cortisone-to-(cortisol + cortisone) ratio was negatively correlated with the amount of BM cortisol (rho = - .914**, *p* < .001) and not associated with BM cortisone (*p* > .05). This suggests a decreased conversion of cortisol to cortisone in initial breast milk. Moreover, we did not find any significant associations between salivary cortisol (AUCg and total decline at T1 or T2), maternal emotional complaints (at T1 and T3), and the BM cortisone-to-(cortisol + cortisone) ratio. All *p*-values were > .05.

### 3.6 Relationship between the BM microbiome and BM glucocorticoids

Kendall’s rank correlations indicated that BM cortisol concentration was significantly associated with BM alpha diversity (τ = 0.1671, FDR = 0.0352*). After adjusting for the FDR [71], BM cortisol was significantly correlated with the family *Staphylococcaceae* (τ = - 0.2047, FDR = 0.0577*) and the genus *Staphylococcus* (τ = - 0.2047, FDR = 0.0885*; see Table 4).

**Table 4.** BM microbiome features in relation to BM cortisol concentrations (*N* = 97 BM samples for associations with microbial taxon abundances, *N* = 96 for alpha and beta diversity measures)

| Category | Item | Variable (T3) | Kendall's tau | p-value | FDR |
| --- | --- | --- | --- | --- | --- |
| Alpha Diversity | Shannon diversity index | Cortisol | 0.0967 | n.s | n.s |
| Alpha Diversity | Richness | Cortisol | 0.1671 | 0.0176* | 0.0352* |

| Category | Item | Variable (T3) | Sum of Squares | Pr(>F) | FDR |
| --- | --- | --- | --- | --- | --- |
| Beta Diversity | Bray-Curtis dissimilarity | Cortisol | 0.4374 | n.s | n.s |
| Beta Diversity | Weighted UniFrac | Cortisol | 0.2497 | n.s | n.s |

| Category | Item | Variable (T3) | Kendall's tau | p-value | FDR |
| --- | --- | --- | --- | --- | --- |
| Family | Gemellaceae | Cortisol | 0.2040 | 0.0038* | 0.0577* |
| Family | Staphylococcaceae | Cortisol | - 0.2047 | 0.0030* | 0.0577* |
| Genus | Gemella | Cortisol | 0.2040 | 0.0038* | 0.0885* |
| Genus | Staphylococcus | Cortisol | - 0.2047 | 0.0030* | 0.0885* |

Note: $p$ -value = statistical significance, \* $p < .05$ ; $Pr(>F) = p$ -value associated with the F statistic; $FDR$ = FDR-adjusted $p$ -value; Cortisol nmol/l = BM cortisol concentration in nmol per litre.

Conversely, no significant associations were found between BM cortisone concentrations and alpha diversity (Shannon diversity index; τ = - 0.0535, FDR = 0.4417, richness; τ = - 0.0595, FDR = 0.4417) or individual BM microbiome taxa (FDR > 0.1). Moreover, both BM cortisol and cortisone showed no significant correlation with the beta diversity of the BM microbiome at T3 (*p* > 0.05).

## 4. Discussion

In this cross-sectional study involving perinatal women, we investigated whether there are any associations between maternal perinatal biological stress, perinatal emotional complaints, and the BM microbiome. Moreover, we analysed psychobiological parameters in relation to BM glucocorticoids and explored the influence of BM glucocorticoid levels on BM bacterial composition. Our main findings suggest that: the BM microbiome is dominated by the genera *Staphylococcus, Streptococcus* and *Gemella;* increased perinatal maternal emotional complaints and salivary AUCg are associated with decreased specific bacterial taxa; and enhanced BM cortisol is linked to alpha diversity and *Staphylococcus*. No associations were found between psychobiological parameters and BM glucocorticoids.

Our results support the view that *Staphylococcus* and *Streptococcus* are predominant bacterial species in BM [77, 78] and reinforce the finding that BM can be an important source of protective bacteria to the infant gut [79]. *Staphylococcus* and *Streptococcus* represented, respectively, 41.1% and 35.2% of the total bacterial isolates in our sample and have been previously identified in stool samples of breast-fed infants [80, 81]. Human milk contains commensal bacteria, mainly non-pathogenic, which compete for nutrients and prevent growth of pathogens by producing acid, hydrogen peroxide or bacteriocins [82]. Conversely, whereas previous studies have identified *Lactobacillus* and *Bifidobacteria* - often considered as part of a healthy microbiota - as common members of BM microbiota [83], we observed very few sequences from these microbes. This difference may be attributable to geographical or environmental differences among the studies, use of colostrum instead of transitional or mature milk, differences in sample processing, analytical techniques, or diet [9].

Studies in both humans and animal models indicate that maternal psychosocial distress is related to negative alterations in gut microbiota [84–87]. This is partially in line with our findings, since breastfeeding may adversely modify infant gut microbiota and BM is recognised as one of the most important elements modulating a child’s gut health [88]. We demonstrated that increased birth-specific anxiety and pregnancy-related psychological symptoms (sum) were associated with a decrease in BM *Alphaproteobacteria*, whereas heightened postpartum anxiety symptoms and postpartum psychological symptoms (sum) were linked to decreased *Alphaproteobacteria, Propionibacteriales, Cutibacterium* and *Pseudomonas*. All of these bacterial communities, although not among the most abundant genera in our BM samples, have been previously reported as members of the human milk or skin microbiota inducing protective responses on the host’s immune system [89, 90]. It was shown that the presence of *Pseudomonas* in BM, alone or in combination with other bacteria, does not define an unhealthy microbial state [91]. This finding suggests that higher maternal emotional complaints contribute to the decrease of commensal, commonly found, gram-positive bacteria in BM. Human milk is in itself a source of commensal bacteria. Commensal strains generally represent ecologically important inhabitants of the human mucosal surfaces [92]. Therefore, this negative alteration in BM microbiota might, at least in part, be driven by increased maternal emotional complaints in pregnancy and postpartum. Importantly, however, not all studies observed this association [90].

Our results revealed that enhanced salivary cortisol, measured by AUCg, was associated with lower *Actinomycetia, Clostridia, Stenotrophomonas, and Kocuria* whereas increased BM cortisol was related to decreased *Staphylococcus*. All these bacteria exert a great impact on regulation of immunophysiological functions, including metabolism and pathogen defense [90, 91]. Here too, we see negative effects of maternal prenatal and postnatal emotional complaints in terms of altering BM microbial composition, and particularly in reducing commensal bacteria. This finding is novel, as prenatal salivary cortisol and BM cortisol have not been previously addressed as potential factors that may influence milk bacterial composition. Notably, we observed a link between higher BM cortisol and increased alpha diversity and richness. Microbiota diversity plays an important role in the maintenance of human immune balance while promoting intestinal immune system maturation [93]. Our finding conflicts with the only previous study to have addressed the effects of postpartum psychosocial stress on BM microbiota, which reported high milk diversity in women with lower psychosocial stress levels [18]. The few other studies assessing the effects of self-reported stress described similar findings in relation to infant gut microbiota diversity [94, 95]. We are the first to publish data specifically targeting BM glucocorticoid effects on BM bacterial composition, thus precluding any direct comparison with previous works. The actual impact of the observed BM alterations remains uncertain, since different taxonomic profiles may retain microbiota functionality due to metabolic plasticity [96]. Future studies examining BM cortisol alterations in BM bacteriome in clinical and non-clinical populations may increase our understanding of the mechanisms linking maternal mental health and BM microbiota.

Finally, we did not find any significant associations between maternal perinatal emotional complaints, salivary cortisol and BM glucocorticoids. Although BM cortisol was previously shown to be strongly correlated with plasma cortisol concentrations and maternal emotional complaints [25, 40], this finding was not replicated in our sample. It is important to acknowledge that some studies also failed to reveal an association between psychological and biological stress during the perinatal phase. For instance, in a single assessment at 28 weeks gestation, Wadhwa and colleagues (1996) [97] found no association between maternal plasma cortisol and stressful life events, perceived stress, or self-reported pregnancy anxiety. Similarly, Shea et al. (2007) [98] examined the cortisol awakening response in pregnant women at 20 weeks gestation and reported no significant associations with self-reported depression or anxiety. Finally, other researchers [99, 100] reported that pregnant women with high stress levels presented with lower cortisol morning levels and a flatter diurnal decline in late pregnancy, with associations between psychological stress and cortisol decline also being observed previously [101]. In contrast, a recent study found that women with higher psychosocial stress had increased levels of BM cortisol [18]. The underlying mechanisms of these opposing effects are unclear, and more studies on glucocorticoid and mineralocorticoid receptor activity are thus needed.

This study is not without limitations. The use of a single sampling time point prevented us from studying intra-individual variability and from exploring changes in BM bacterial profiles over time. Moreover, the high sensitivity of the next-generation sequencing used in our analyses can detect minimal amounts of contaminant DNA, especially in samples carrying a low bacterial load such as BM, making it difficult to characterise BM microbes without the influence of other DNA sources (e.g. skin). Strengths of our study include our large sample size, the consistent methods for sample collection, processing and sequencing, and the robust statistical analysis.

In sum, we provide initial insights into the dynamics of how maternal psychobiological aspects in pregnancy and postpartum may affect BM composition. We report that increased perinatal emotional complaints and prenatal salivary cortisol AUCg are associated with decreased specific commensal bacteria, whereas enhanced BM cortisol is linked to higher BM alpha diversity and reduced *Staphylococcus*. No associations were found between psychobiological parameters and BM glucocorticoids. Future studies exploring the consequence of these alterations on the infant gut and development are warranted.

## Abbreviations

ASV: amplicon sequence variant
AUCg: area under the curve with respect to the ground
BH: Benjamini-Hochberg
BM: Breast milk
dbRDA: distance-based redundancy analysis
EPDS: Edinburgh Postnatal Depression Scale
FDR: False discovery rate
GAS: Geburts-Angst-Skala
GHQ-12: General Health Questionnaire
HPA axis: hypothalamic-pituitary-adrenal axis
NuPDQ: Prenatal Distress Questionnaire
PSS-10: Perceived Stress Scale
STAI-S: State-Trate Anxiety Inventory, state scale

## Acknowledgements

We wish to express our gratitude to all women who participated in this study. We are grateful to our study team members Annika Schneider, Eileen Stephan, Nathalie Appenzeller, Rhea Sarah Gesù, Tamara Grossrieder, Tamara Lovrinovic, Seraina Weber, Stéphanie Loeffel, Simon Hühni and Firouzeh Farahmand for their research assistantship. Special thanks to Rausch AG, Medela Schweiz AG, MAM Baby AG and Bimbosan that freely provided gift sets for the study participants. Finally, we would like to thank the Swiss Federal Food Safety and Veterinary Office as well as the Research Group for Food Perception from the ZHAW School of Life Sciences for their support.

## Authors’ Contributions

RAC and UE conceived the presented idea. ND and RAC conducted research. ND analysed the data. RAC and UE verified the analytical methods and contributed to the interpretation of the results. RAC supervised the manuscript. ND took the lead in writing the manuscript. All authors provided critical feedback and contributed to the final version of the manuscript.

## References

[1] E. J. Mayer-Davis, S. L. Rifas-Shiman, L. Zhou, F. B. Hu, G. A. Colditz, and M. W. Gillman, “Breast-feeding and risk for childhood obesity: does maternal diabetes or obesity status matter?,” Diabetes care, vol. 29, no. 10, pp. 2231–2237, 2006, doi: 10.2337/dc06-0974.

[2] C. G. Victora et al., “Breastfeeding in the 21st century: epidemiology, mechanisms, and lifelong effect,” The Lancet, vol. 387, no. 10017, pp. 475–490, 2016, doi: 10.1016/S0140-6736(15)01024-7.

[3] T. M. Samuel, Q. Zhou, F. Giuffrida, D. Munblit, V. Verhasselt, and S. K. Thakkar, “Nutritional and Non-nutritional Composition of Human Milk Is Modulated by Maternal, Infant, and Methodological Factors,” Frontiers in nutrition, vol. 7, p. 576133, 2020, doi: 10.3389/fnut.2020.576133.

[4] O. Ballard and A. L. Morrow, “Human milk composition: nutrients and bioactive factors,” Pediatric clinics of North America, vol. 60, no. 1, pp. 49–74, 2013, doi: 10.1016/j.pcl.2012.10.002.

[5] V. Notarbartolo, M. Giuffrè, C. Montante, G. Corsello, and M. Carta, “Composition of Human Breast Milk Microbiota and Its Role in Children’s Health,” Pediatric gastroenterology, hepatology & nutrition, vol. 25, no. 3, pp. 194–210, 2022, doi: 10.5223/pghn.2022.25.3.194.

[6] A. Togo, J.-C. Dufour, J.-C. Lagier, G. Dubourg, D. Raoult, and M. Million, “Repertoire of human breast and milk microbiota: a systematic review,” Future microbiology, vol. 14, pp. 623–641, 2019, doi: 10.2217/fmb-2018-0317.

[7] C. A. Moubareck, “Human Milk Microbiota and Oligosaccharides: A Glimpse into Benefits, Diversity, and Correlations,” Nutrients, vol. 13, no. 4, 2021, doi: 10.3390/nu13041123.

[8] A. Diez-Sampedro, M. Flowers, M. Olenick, T. Maltseva, and G. Valdes, “Women’s Choice Regarding Breastfeeding and Its Effect on Well-Being,” Nursing for women’s health, vol. 23, no. 5, pp. 383–389, 2019, doi: 10.1016/j.nwh.2019.08.002.

[9] K. M. Hunt et al., “Characterization of the diversity and temporal stability of bacterial communities in human milk,” PloS one, vol. 6, no. 6, e21313, 2011, doi: 10.1371/journal.pone.0021313.

[10] M. Kortesniemi et al., “Human milk metabolome is associated with symptoms of maternal psychological distress and milk cortisol,” Food chemistry, vol. 356, p. 129628, 2021, doi: 10.1016/j.foodchem.2021.129628.

[11] M. G. Di Benedetto, C. Bottanelli, A. Cattaneo, C. M. Pariante, and A. Borsini, “Nutritional and immunological factors in breast milk: A role in the intergenerational transmission from maternal psychopathology to child development,” Brain, behavior, and immunity, vol. 85, pp. 57–68, 2020, doi: 10.1016/j.bbi.2019.05.032.

[12] H. Demmelmair, E. Jiménez, M. C. Collado, S. Salminen, and M. K. McGuire, “Maternal and Perinatal Factors Associated with the Human Milk Microbiome,” Current developments in nutrition, vol. 4, no. 4, nzaa027, 2020, doi: 10.1093/cdn/nzaa027.

[13] R. Enaud et al., “The Gut-Lung Axis in Health and Respiratory Diseases: A Place for Inter-Organ and Inter-Kingdom Crosstalks,” Frontiers in cellular and infection microbiology, vol. 10, p. 9, 2020, doi: 10.3389/fcimb.2020.00009.

[14] S. K. Dogra, J. Doré, and S. Damak, “Gut Microbiota Resilience: Definition, Link to Health and Strategies for Intervention,” Frontiers in microbiology, vol. 11, p. 572921, 2020, doi: 10.3389/fmicb.2020.572921.

[15] K. Hufnagl, I. Pali-Schöll, F. Roth-Walter, and E. Jensen-Jarolim, “Dysbiosis of the gut and lung microbiome has a role in asthma,” Seminars in immunopathology, vol. 42, no. 1, pp. 75–93, 2020, doi: 10.1007/s00281-019-00775-y.

[16] K. G. Margolis, J. F. Cryan, and E. A. Mayer, “The Microbiota-Gut-Brain Axis: From Motility to Mood,” Gastroenterology, vol. 160, no. 5, pp. 1486–1501, 2021, doi: 10.1053/j.gastro.2020.10.066.

[17] G. Oikonomou et al., “Milk Microbiota: What Are We Exactly Talking About?,” Frontiers in microbiology, vol. 11, p. 60, 2020, doi: 10.3389/fmicb.2020.00060.

[18] P. D. Browne et al., “Human Milk Microbiome and Maternal Postnatal Psychosocial Distress,” Frontiers in microbiology, vol. 10, p. 2333, 2019, doi: 10.3389/fmicb.2019.02333.

[19] M. W. Groer, M. W. Davis, and J. Hemphill, “Postpartum stress: current concepts and the possible protective role of breastfeeding,” Journal of obstetric, gynecologic, and neonatal nursing : JOGNN, vol. 31, no. 4, pp. 411–417, 2002, doi: 10.1111/j.1552-6909.2002.tb00063.x.

[20] R. J. Ruiz and J. T. Fullerton, “The measurement of stress in pregnancy,” Nursing & health sciences, vol. 1, no. 1, pp. 19–25, 1999, doi: 10.1046/j.1442-2018.1999.00004.x.

[21] R. H. van den Bergh et al., “Prenatal developmental origins of behavior and mental health: The influence of maternal stress in pregnancy,” Neuroscience and biobehavioral reviews, vol. 117, pp. 26–64, 2020, doi: 10.1016/j.neubiorev.2017.07.003.

[22] S. M. Woods, J. L. Melville, Y. Guo, M.-Y. Fan, and A. Gavin, “Psychosocial stress during pregnancy,” American journal of obstetrics and gynecology, vol. 202, no. 1, 61.e1-7, 2010, doi: 10.1016/j.ajog.2009.07.041.

[23] A. Kawano, Y. Emori, and S. Miyagawa, “Association between stress-related substances in saliva and immune substances in breast milk in puerperae,” Biological research for nursing, vol. 10, no. 4, pp. 350–355, 2009, doi: 10.1177/1099800409331892.

[24] S. Pundir et al., “Maternal influences on the glucocorticoid concentrations of human milk: The STEPS study,” Clinical nutrition (Edinburgh, Scotland), vol. 38, no. 4, pp. 1913–1920, 2019, doi: 10.1016/j.clnu.2018.06.980.

[25] J. J. Hollanders, A. C. Heijboer, B. van der Voorn, J. Rotteveel, and M. J. J. Finken, “Nutritional programming by glucocorticoids in breast milk: Targets, mechanisms and possible implications,” Best practice & research. Clinical endocrinology & metabolism, vol. 31, no. 4, pp. 397–408, 2017, doi: 10.1016/j.beem.2017.10.001.

[26] A. Stein et al., “Effects of perinatal mental disorders on the fetus and child,” The Lancet, vol. 384, no. 9956, pp. 1800–1819, 2014, doi: 10.1016/S0140-6736(14)61277-0.

[27] K. M. Sawyer, P. A. Zunszain, P. Dazzan, and C. M. Pariante, “Intergenerational transmission of depression: clinical observations and molecular mechanisms,” Molecular psychiatry, vol. 24, no. 8, pp. 1157–1177, 2019, doi: 10.1038/s41380-018-0265-4.

[28] L. Drago, M. Toscano, R. de Grandi, E. Grossi, E. M. Padovani, and D. G. Peroni, “Microbiota network and mathematic microbe mutualism in colostrum and mature milk collected in two different geographic areas: Italy versus Burundi,” The ISME journal, vol. 11, no. 4, pp. 875–884, 2017, doi: 10.1038/ismej.2016.183.

[29] L. Fernández et al., “The human milk microbiota: origin and potential roles in health and disease,” Pharmacological research, vol. 69, no. 1, pp. 1–10, 2013, doi: 10.1016/j.phrs.2012.09.001.

[30] A. J. Macpherson and T. Uhr, “Induction of protective IgA by intestinal dendritic cells carrying commensal bacteria,” Science (New York, N.Y.), vol. 303, no. 5664, pp. 1662– 1665, 2004, doi: 10.1126/science.1091334.

[31] P. F. Perez et al., “Bacterial imprinting of the neonatal immune system: lessons from maternal cells?,” Pediatrics, vol. 119, no. 3, e724–32, 2007, doi: 10.1542/peds.2006-1649.

[32] L. F. Stinson et al., “The human milk microbiome: who, what, when, where, why, and how?,” Nutrition reviews, vol. 79, no. 5, pp. 529–543, 2021, doi: 10.1093/nutrit/nuaa029.

[33] E. A. Mayer and E. Y. Hsiao, “The Gut and Its Microbiome as Related to Central Nervous System Functioning and Psychological Well-being: Introduction to the Special Issue of Psychosomatic Medicine,” Psychosomatic medicine, vol. 79, no. 8, pp. 844–846, 2017, doi: 10.1097/PSY.0000000000000525.

[34] G. de Palma, S. M. Collins, P. Bercik, and E. F. Verdu, “The microbiota-gut-brain axis in gastrointestinal disorders: stressed bugs, stressed brain or both?,” The Journal of physiology, vol. 592, no. 14, pp. 2989–2997, 2014, doi: 10.1113/jphysiol.2014.273995.

[35] V. Osadchiy, C. R. Martin, and E. A. Mayer, “The Gut-Brain Axis and the Microbiome: Mechanisms and Clinical Implications,” Clinical gastroenterology and hepatology : the official clinical practice journal of the American Gastroenterological Association, vol. 17, no. 2, pp. 322–332, 2019, doi: 10.1016/j.cgh.2018.10.002.

[36] B. Misiak et al., “The HPA axis dysregulation in severe mental illness: Can we shift the blame to gut microbiota?,” Progress in neuro-psychopharmacology & biological psychiatry, vol. 102, p. 109951, 2020, doi: 10.1016/j.pnpbp.2020.109951.

[37] A. Harris and J. Seckl, “Glucocorticoids, prenatal stress and the programming of disease,” Hormones and behavior, vol. 59, no. 3, pp. 279–289, 2011, doi: 10.1016/j.yhbeh.2010.06.007.

[38] G. A. Kaltsas and G. P. Chrousos, Eds., The neuroendocrinology of stress, 3rd ed. Cambridge, UK: Cambridge University Press, 2007.

[39] R. M. Sapolsky, L. M. Romero, and A. U. Munck, “How do glucocorticoids influence stress responses? Integrating permissive, suppressive, stimulatory, and preparative actions,” Endocr Rev, vol. 21, no. 1, pp. 55–89, 2000, doi: 10.1210/edrv.21.1.0389.

[40] F. R. Patacchioli et al., “Maternal plasma and milk free cortisol during the first 3 days of breast-feeding following spontaneous delivery or elective cesarean section,” Gynecologic and obstetric investigation, vol. 34, no. 3, pp. 159–163, 1992, doi: 10.1159/000292751.

[41] M. Aparicio et al., “Human milk cortisol and immune factors over the first three postnatal months: Relations to maternal psychosocial distress,” PloS one, vol. 15, no. 5, e0233554, 2020, doi: 10.1371/journal.pone.0233554.

[42] S. Hart, L. Boylan, B. Border, S. R. Carroll, D. McGunegle, and R. M. Lampe, “Breast milk levels of cortisol and Secretory Immunoglobulin A (SIgA) differ with maternal mood and infant neuro-behavioral functioning,” Infant Behavior and Development, vol. 27, no. 1, pp. 101–106, 2004, doi: 10.1016/j.infbeh.2003.06.002.

[43] M. Romijn et al., “The Association between Maternal Stress and Glucocorticoid Rhythmicity in Human Milk,” Nutrients, vol. 13, no. 5, 2021, doi: 10.3390/nu13051608.

[44] H. A. Tucker and J. W. Schwalm, “Glucocorticoids in mammary tissue and milk,” Journal of animal science, vol. 45, no. 3, pp. 627–634, 1977, doi: 10.2527/jas1977.453627x.

[45] M. Klein et al., “The German version of the Perceived Stress Scale - psychometric characteristics in a representative German community sample,” BMC psychiatry, vol. 16, p. 159, 2016, doi: 10.1186/s12888-016-0875-9.

[46] S. Cohen, T. Kamarck, and R. Mermelstein, “A Global Measure of Perceived Stress,” Journal of Health and Social Behavior, vol. 24, no. 4, p. 385, 1983, doi: 10.2307/2136404.

[47] N. Schmitz, J. Kruse, and W. Tress, “Psychometric properties of the General Health Questionnaire (GHQ-12) in a German primary care sample,” Acta psychiatrica Scandinavica, vol. 100, no. 6, pp. 462–468, 1999, doi: 10.1111/j.1600-0447.1999.tb10898.x.

[48] M. Romppel, E. Braehler, M. Roth, and H. Glaesmer, “What is the General Health Questionnaire-12 assessing? Dimensionality and psychometric properties of the General Health Questionnaire-12 in a large scale German population sample,” Comprehensive psychiatry, vol. 54, no. 4, pp. 406–413, 2013, doi: 10.1016/j.comppsych.2012.10.010.

[49] P. Goldberg and B. Blackwell, “Psychiatric illness in general practice. A detailed study using a new method of case identification,” British medical journal, vol. 1, no. 5707, pp. 439–443, 1970, doi: 10.1136/bmj.2.5707.439.

[50] M. Hankins, “The factor structure of the twelve item General Health Questionnaire (GHQ-12): the result of negative phrasing?,” Clinical practice and epidemiology in mental health : CP & EMH, vol. 4, p. 10, 2008, doi: 10.1186/1745-0179-4-10.

[51] S. M. Ibrahim and M. Lobel, “Conceptualization, measurement, and effects of pregnancy-specific stress: review of research using the original and revised Prenatal Distress Questionnaire,” Journal of behavioral medicine, vol. 43, no. 1, pp. 16–33, 2020, doi: 10.1007/s10865-019-00068-7.

[52] M. Lobel, D. L. Cannella, J. E. Graham, C. DeVincent, J. Schneider, and B. A. Meyer, “Pregnancy-specific stress, prenatal health behaviors, and birth outcomes,” Health psychology : official journal of the Division of Health Psychology, American Psychological Association, vol. 27, no. 5, pp. 604–615, 2008, doi: 10.1037/a0013242.

[53] A. M. Bergant, T. Nguyen, K. Heim, H. Ulmer, and O. Dapunt, “Deutschsprachige Fassung und Validierung der “Edinburgh postnatal depression scale”,” (in ger), Deutsche medizinische Wochenschrift (1946), vol. 123, no. 3, pp. 35–40, 1998, doi: 10.1055/s-2007-1023895.

[54] H. Lukesch, Geburts-Angst-Skala G-A-S. Göttingen: Hogrefe, 1983.

[55] A. Bratsikas, “Assessment of psychobiological stress reactivity and its relation to postpartum mood states,” University of Zurich, 2006.

[56] L. Laux, Das State-Trait-Angstinventar (STAI) : theoretische Grundlagen und Handanweisung: Beltz, 1981. [Online]. Available: https://fis.uni-bamberg.de/handle/uniba/26756

[57] C. Spielberger and S. Sydeman, State-trait anxiety inventory and state-trait anger expression inventory. The use of psychological testing for treatment planning and outcome assessment, 1994. [Online]. Available: https://scholar.google.com/citations?user=l5bq0t4aaaaj&hl=en&oi=sra

[58] Spielberger C. D., “Manual for the State-trait Anxietry, Inventory,” Consulting Psychologist, 1970. [Online]. Available: https://cir.nii.ac.jp/crid/1572261549249149184

[59] Y. Gidron, “Trait Anxiety,” in Encyclopedia of Behavioral Medicine, M. Gellman and J. R. Turner, Eds., New York, NY: Springer New York, 2016, pp. 1–2.

[60] J. Heron, T. G. O’Connor, J. Evans, J. Golding, and V. Glover, “The course of anxiety and depression through pregnancy and the postpartum in a community sample,” Journal of affective disorders, vol. 80, no. 1, pp. 65–73, 2004, doi: 10.1016/j.jad.2003.08.004.

[61] I. Nast, M. Bolten, G. Meinlschmidt, and D. H. Hellhammer, “How to measure prenatal stress? A systematic review of psychometric instruments to assess psychosocial stress during pregnancy,” Paediatric and perinatal epidemiology, vol. 27, no. 4, pp. 313–322, 2013, doi: 10.1111/ppe.12051.

[62] B. van der Voorn, M. de Waard, J. B. van Goudoever, J. Rotteveel, A. C. Heijboer, and M. J. Finken, “Breast-Milk Cortisol and Cortisone Concentrations Follow the Diurnal Rhythm of Maternal Hypothalamus-Pituitary-Adrenal Axis Activity,” The Journal of nutrition, vol. 146, no. 11, pp. 2174–2179, 2016, doi: 10.3945/jn.116.236349.

[63] M. K. McGuire et al., “What’s normal? Oligosaccharide concentrations and profiles in milk produced by healthy women vary geographically,” The American journal of clinical nutrition, vol. 105, no. 5, pp. 1086–1100, 2017, doi: 10.3945/ajcn.116.139980.

[64] B. van der Voorn et al., “Determination of cortisol and cortisone in human mother’s milk,” Clinica chimica acta; international journal of clinical chemistry, vol. 444, pp. 154–155, 2015, doi: 10.1016/j.cca.2015.02.015.

[65] A. Klindworth et al., “Evaluation of general 16S ribosomal RNA gene PCR primers for classical and next-generation sequencing-based diversity studies,” Nucleic acids research, vol. 41, no. 1, e1, 2013, doi: 10.1093/nar/gks808.

[66] F. Faul, E. Erdfelder, A. Buchner, and A.-G. Lang, “Statistical power analyses using G*Power 3.1: tests for correlation and regression analyses,” Behavior research methods, vol. 41, no. 4, pp. 1149–1160, 2009, doi: 10.3758/BRM.41.4.1149.

[67] C. Pruessner, C. Kirschbaum, G. Meinlschmid, and D. H. Hellhammer, “Two formulas for computation of the area under the curve represent measures of total hormone concentration versus time-dependent change,” Psychoneuroendocrinology, vol. 28, no. 7, pp. 916–931, 2003, doi: 10.1016/S0306-4530(02)00108-7.

[68] P. La Marca-Ghaemmaghami, R. La Marca, S. M. Dainese, M. Haller, R. Zimmermann, and U. Ehlert, “The association between perceived emotional support, maternal mood, salivary cortisol, salivary cortisone, and the ratio between the two compounds in response to acute stress in second trimester pregnant women,” Journal of psychosomatic research, vol. 75, no. 4, pp. 314–320, 2013, doi: 10.1016/j.jpsychores.2013.08.010.

[69] R. Benediktsson, A. A. Calder, C. R. Edwards, and J. R. Seckl, “Placental 11 beta-hydroxysteroid dehydrogenase: a key regulator of fetal glucocorticoid exposure,” Clinical endocrinology, vol. 46, no. 2, pp. 161–166, 1997, doi: 10.1046/j.1365-2265.1997.1230939.x.

[70] B. J. Callahan, P. J. McMurdie, M. J. Rosen, A. W. Han, A. J. A. Johnson, and S. P. Holmes, “DADA2: High-resolution sample inference from Illumina amplicon data,” Nature methods, vol. 13, no. 7, pp. 581–583, 2016, doi: 10.1038/nmeth.3869.

[71] Y. Benjamini and Y. Hochberg, “Controlling the False Discovery Rate: A Practical and Powerful Approach to Multiple Testing,” Journal of the Royal Statistical Society: Series B (Methodological), vol. 57, no. 1, pp. 289–300, 1995, doi: 10.1111/j.2517-6161.1995.tb02031.x.

[72] G. Falony et al., “Population-level analysis of gut microbiome variation,” Science (New York, N.Y.), vol. 352, no. 6285, pp. 560–564, 2016, doi: 10.1126/science.aad3503.

[73] E. Bayar, P. R. Bennett, D. Chan, L. Sykes, and D. A. MacIntyre, “The pregnancy microbiome and preterm birth,” Seminars in immunopathology, vol. 42, no. 4, pp. 487– 499, 2020, doi: 10.1007/s00281-020-00817-w.

[74] S. L. Gillespie, A. M. Mitchell, J. M. Kowalsky, and L. M. Christian, “Maternal parity and perinatal cortisol adaptation: The role of pregnancy-specific distress and implications for postpartum mood,” Psychoneuroendocrinology, vol. 97, pp. 86–93, 2018, doi: 10.1016/j.psyneuen.2018.07.008.

[75] C. Gomez-Gallego, I. Garcia-Mantrana, S. Salminen, and M. C. Collado, “The human milk microbiome and factors influencing its composition and activity,” Seminars in fetal & neonatal medicine, vol. 21, no. 6, pp. 400–405, 2016, doi: 10.1016/j.siny.2016.05.003.

[76] N. T. Mueller et al., “Birth mode-dependent association between pre-pregnancy maternal weight status and the neonatal intestinal microbiome,” Scientific reports, vol. 6, p. 23133, 2016, doi: 10.1038/srep23133.

[77] L. Carroll, “BACTERIOLOGICAL CRITERIA FOR FEEDING RAW BREAST-MILK TO BABIES ON NEONATAL UNITS,” The Lancet, vol. 314, no. 8145, pp. 732– 733, 1979, doi: 10.1016/s0140-6736(79)90654-8.

[78] A. I. Eidelman and G. Szilagyi, “Patterns of bacterial colonization of human milk,” Obstetrics and gynecology, vol. 53, no. 5, pp. 550–552, 1979.

[79] M. P. Heikkilä and P. E. J. Saris, “Inhibition of Staphylococcus aureus by the commensal bacteria of human milk,” Journal of applied microbiology, vol. 95, no. 3, pp. 471–478, 2003, doi: 10.1046/j.1365-2672.2003.02002.x.

[80] P. V. Kirjavainen, E. Apostolou, T. Arvola, S. J. Salminen, G. R. Gibson, and E. Isolauri, “Characterizing the composition of intestinal microflora as a prospective treatment target in infant allergic disease,” FEMS immunology and medical microbiology, vol. 32, no. 1, pp. 1–7, 2001, doi: 10.1111/j.1574-695X.2001.tb00526.x.

[81] C. F. Favier, E. E. Vaughan, W. M. de Vos, and A. D. L. Akkermans, “Molecular monitoring of succession of bacterial communities in human neonates,” Applied and environmental microbiology, vol. 68, no. 1, pp. 219–226, 2002, doi: 10.1128/AEM.68.1.219-226.2002.

[82] P. Huovinen, “Bacteriotherapy: the time has come : Bacterial interference is an increasingly attractive approach to prevention and therapy,” BMJ : British Medical Journal, vol. 323, no. 7309, pp. 353–354, 2001.

[83] M. C. Collado, S. Delgado, A. Maldonado, and J. M. Rodríguez, “Assessment of the bacterial diversity of breast milk of healthy women by quantitative real-time PCR,” Letters in applied microbiology, vol. 48, no. 5, pp. 523–528, 2009, doi: 10.1111/j.1472-765X.2009.02567.x.

[84] J. F. Cryan and T. G. Dinan, “Mind-altering microorganisms: the impact of the gut microbiota on brain and behaviour,” Nature reviews. Neuroscience, vol. 13, no. 10, pp. 701–712, 2012, doi: 10.1038/nrn3346.

[85] J. D. Galley, N. M. Parry, B. M. M. Ahmer, J. G. Fox, and M. T. Bailey, “The commensal microbiota exacerbate infectious colitis in stressor-exposed mice,” Brain, behavior, and immunity, vol. 60, pp. 44–50, 2017, doi: 10.1016/j.bbi.2016.09.010.

[86] J. D. Galley et al., “Exposure to a social stressor disrupts the community structure of the colonic mucosa-associated microbiota,” BMC microbiology, vol. 14, p. 189, 2014, doi: 10.1186/1471-2180-14-189.

[87] C. Hechler et al., “Association between Psychosocial Stress and Fecal Microbiota in Pregnant Women,” Scientific reports, vol. 9, no. 1, p. 4463, 2019, doi: 10.1038/s41598-019-40434-8.

[88] R. Cabrera-Rubio, M. C. Collado, K. Laitinen, S. Salminen, E. Isolauri, and A. Mira, “The human milk microbiome changes over lactation and is shaped by maternal weight and mode of delivery,” The American journal of clinical nutrition, vol. 96, no. 3, pp. 544–551, 2012, doi: 10.3945/ajcn.112.037382.

[89] L. Ruiz et al., “Comparison of Two Approaches for the Metataxonomic Analysis of the Human Milk Microbiome,” Frontiers in cellular and infection microbiology, vol. 11, p. 622550, 2021, doi: 10.3389/fcimb.2021.622550.

[90] A. Ojo-Okunola et al., “Influence of Socio-Economic and Psychosocial Profiles on the Human Breast Milk Bacteriome of South African Women,” Nutrients, vol. 11, no. 6, 2019, doi: 10.3390/nu11061390.

[91] R. W. Hyman, M. Fukushima, L. Diamond, J. Kumm, L. C. Giudice, and R. W. Davis, “Microbes on the human vaginal epithelium,” Proceedings of the National Academy of Sciences of the United States of America, vol. 102, no. 22, pp. 7952–7957, 2005, doi: 10.1073/pnas.0503236102.

[92] L. Grozdanov et al., “Analysis of the genome structure of the nonpathogenic probiotic Escherichia coli strain Nissle 1917,” Journal of bacteriology, vol. 186, no. 16, pp. 5432– 5441, 2004, doi: 10.1128/JB.186.16.5432-5441.2004.

[93] M. Selma-Royo, J. Calvo Lerma, E. Cortés-Macías, and M. C. Collado, “Human milk microbiome: From actual knowledge to future perspective,” Seminars in perinatology, vol. 45, no. 6, p. 151450, 2021, doi: 10.1016/j.semperi.2021.151450.

[94] S. R. Knowles, E. A. Nelson, and E. A. Palombo, “Investigating the role of perceived stress on bacterial flora activity and salivary cortisol secretion: a possible mechanism underlying susceptibility to illness,” Biological psychology, vol. 77, no. 2, pp. 132–137, 2008, doi: 10.1016/j.biopsycho.2007.09.010.

[95] A. Kato-Kataoka et al., “Fermented Milk Containing Lactobacillus casei Strain Shirota Preserves the Diversity of the Gut Microbiota and Relieves Abdominal Dysfunction in Healthy Medical Students Exposed to Academic Stress,” Applied and environmental microbiology, vol. 82, no. 12, pp. 3649–3658, 2016, doi: 10.1128/AEM.04134-15.

[96] A. Moya and M. Ferrer, “Functional Redundancy-Induced Stability of Gut Microbiota Subjected to Disturbance,” Trends in microbiology, vol. 24, no. 5, pp. 402–413, 2016, doi: 10.1016/j.tim.2016.02.002.

[97] P. D. Wadhwa, C. Dunkel-Schetter, A. Chicz-DeMet, M. Porto, and C. A. Sandman, “Prenatal psychosocial factors and the neuroendocrine axis in human pregnancy,” Psychosomatic medicine, vol. 58, no. 5, pp. 432–446, 1996, doi: 10.1097/00006842-199609000-00006.

[98] A. K. Shea, D. L. Streiner, A. Fleming, M. V. Kamath, K. Broad, and M. Steiner, “The effect of depression, anxiety and early life trauma on the cortisol awakening response during pregnancy: preliminary results,” Psychoneuroendocrinology, vol. 32, 8-10, pp. 1013–1020, 2007, doi: 10.1016/j.psyneuen.2007.07.006.

[99] S. F. Suglia, J. Staudenmayer, S. Cohen, M. B. Enlow, J. W. Rich-Edwards, and R. J. Wright, “Cumulative Stress and Cortisol Disruption among Black and Hispanic Pregnant Women in an Urban Cohort,” Psychological trauma : theory, research, practice and policy, vol. 2, no. 4, pp. 326–334, 2010, doi: 10.1037/a0018953.

[100] T. G. O’Connor, W. Tang, M. A. Gilchrist, J. A. Moynihan, E. K. Pressman, and E. R. Blackmore, “Diurnal cortisol patterns and psychiatric symptoms in pregnancy: short-term longitudinal study,” Biological psychology, vol. 96, pp. 35–41, 2014, doi: 10.1016/j.biopsycho.2013.11.002.

[101] G. Obel, M. Hedegaard, T. B. Henriksen, N. J. Secher, J. Olsen, and S. Levine, “Stress and salivary cortisol during pregnancy,” Psychoneuroendocrinology, vol. 30, no. 7, pp. 647–656, 2005, doi: 10.1016/j.psyneuen.2004.11.006.

